# Heritable variation in bleaching responses and its functional genomic basis in reef-building corals (*Orbicella faveolata*)

**DOI:** 10.1101/185595

**Authors:** Katherine E. Dziedzic, Holland Elder, Hannah Tavalire, Eli Meyer

**Affiliations:** Department of Integrative Biology, Oregon State University, Corvallis OR; Institute of Ecology and Evolution, University of Oregon, Eugene OR

**Keywords:** heritability, genomic, thermal tolerance, coral bleaching

## Abstract

Reef-building corals are highly sensitive to rising ocean temperatures, and substantial adaptation will be required for corals and the ecosystems they support to persist in changing ocean conditions. Genetic variation that might support adaptive responses has been measured in larval stages of some corals, but these estimates remain unavailable for adult corals and the functional basis of this variation remains unclear. In this study, we focused on the potential for adaptation in *Orbicella faveolata*, a dominant reef-builder in the Caribbean. We conducted thermal stress experiments using corals collected from natural populations in Bocas del Toro, Panama, and used multilocus SNP genotypes to estimate genetic relatedness among samples. This allowed us to estimate narrow-sense heritability of variation in bleaching responses, revealing that variation in these responses was highly heritable (*h^2^*=0.58). This suggests substantial potential for adaptive responses to warming by natural populations of *O. faveolata* in this region. We further investigated the functional basis for this variation using genomic and transcriptomic approaches. We used a publicly available genetic linkage map and genome assembly to map markers associated with bleaching responses, identifying twelve markers associated with variation in bleaching responses. We also profiled gene expression in corals with contrasting bleaching phenotypes, uncovering substantial differences in transcriptional stress responses between heat-tolerant and heat-susceptible corals. Together, our findings contribute to the growing body of evidence that natural populations of corals possess genetic variation in thermal stress responses that may potentially support adaptive responses to rising ocean temperatures.

## Introduction

Coral reefs are one of the most diverse and complex ecosystems in the world. They provide habitat for hundreds of thousands of invertebrates and fish, protect coastal environments, and support a variety of resources for local communities. Unfortunately, the invaluable ecosystem services they provide are at risk of being lost as coral reefs worldwide continue to decline. Coral reefs are particularly sensitive to increases in sea surface temperature and have undergone worldwide degradation as the oceans have warmed (Brown, 1997; Hoegh-Guldberg & Jones, 1999; Baker *et al.*, 2004; Eakin *et al.*, 2009). Bleaching events, which reflect the breakdown of symbiotic relationships between corals and dinoflagellates (*Symbiodiniaceae*, formerly *Symbiodinium spp.* (LaJeunesse *et al.*, 2018)) resulting from environmental stress, have increased in frequency and severity over the past few decades (Hughes, 2003; Donner *et al.*, 2005; Hoegh-Guldberg *et al.*, 2007). In the past three years alone, 30-50% of coral reefs have declined in some areas along the Great Barrier Reef (Hughes *et al.*, 2017). This dramatic decline in such a short period of time demands an increased understanding of the potential for these ecosystems to persist into the future.

In order to persist, corals will need to increase their thermal tolerance to cope with ocean warming. It is already known that coral species have differing thermal capacities due to extrinsic factors such as variation in their environment, as well as intrinsic mechanisms to deal with acute and long-term stress events, such as varying associations in symbiont type (Baker *et al.*, 2004; Van Oppen *et al.*, 2005; Jones *et al.*, 2008) or changes in gene expression in the coral host (Bellantuono *et al.*, 2012a; Kenkel *et al.*, 2013). Importantly, bleaching thresholds for some species have been shown to change over time (Fitt *et al.*, 2001; Grottoli *et al.*, 2014). Models that consider both environmental conditions and these changing thresholds suggest that the fate of corals during the next century may be strongly affected by long-term adaptive changes in bleaching thresholds (Donner *et al.*, 2005; D’Angelo *et al.*, 2015). Changes in bleaching thresholds may occur in populations through adaptation (Meyer *et al.*, 2009; Coles & Riegl, 2013; Palumbi *et al.*, 2014), or in individual corals through acclimatization (Jones & Berkelmans, 2010; Oliver & Palumbi, 2011).

Adaptation through genetic change can play a large role in allowing populations to persist in a changing environment. Genetic variation within populations in fitness-related traits, including resistance to environmental stress, supports adaptive responses to selection (Falconer & Mackay, 1996; Barrett & Schluter, 2008). Predicting the adaptive potential of a trait requires an understanding of the proportion of phenotypic variation resulting from genetic factors (Falconer and Mackay, 1996). However, the relative contributions of environmental and genetic factors to variation in thermal tolerance of corals remain poorly understood (Császár *et al.*, 2010). Some studies have provided evidence for corals’ adaptive capacity, demonstrating thermal tolerance differences between local populations (Palumbi *et al.*, 2014; Howells *et al.*, 2016) and considerable heritable variation in thermal tolerance in coral larvae (Dixon *et al.*, 2015) and algal symbionts (Császár *et al.*, 2010). These examples have provided an important first demonstration that genetic potential for adaptation exists in natural populations, but many questions still remain.

Global sea surface temperatures are predicted to rise 1-2°C by the end of the century, and thermally sensitive organisms like reef-building corals will require substantial adaptive responses. Adaptive responses to selection depend on the change in a population’s phenotypic mean and the narrow-sense heritability (*h^2^*), the proportion of total phenotypic variance that is due to additive genetic factors (Falconer and Mackay, 1996). Quantitative estimates of this parameter allow us to estimate the expected evolutionary change in a trait per generation (Visscher *et al.*, 2008; Morrissey *et al.*, 2012). In order to estimate selection responses in corals and consider rates of adaptation, we need to quantify heritability in thermal tolerance. Currently, very few studies provide heritability estimates for coral species and their algal symbionts, particularly in natural populations (Meyer *et al.*, 2009b; Dixon *et al.*, 2015; Kenkel *et al.*, 2015). Previous studies have focused on larval stages for important advantages in experimental design, leaving it unclear whether the high heritabilities estimated in larval responses to elevated temperatures (Meyer *et al.*, 2009, Dixon *et al.*, 2015) can be generalized to understand responses to selection on the adult stage. Further, since the heritability of a trait is specific to a particular population and environment in which it is measured, it remains unclear whether previous estimates of *h^2^* from Indo-Pacific Acroporids can be generalized to evaluate adaptive potential in other regions and species.

The Caribbean has seen dramatic reductions in coral cover over the last thirty years (Hughes & Tanner, 2000; Gardner, 2003) and the potential for existing populations to recover or adapt to changing ocean conditions remains unknown. To understand the potential for adaptation by corals in this region, we investigated the mechanisms that may enable long-term adaptation by investigating heritable variation in thermal tolerance and its genomic basis in *Orbicella faveolata*, a dominant reef-builder in the Caribbean. Our studies aim to quantify the contribution of genetic factors to variation in thermal tolerance of corals, and identify genetic markers and genes associated with this variation. Together our findings providing new insights into the potential for adaptive changes in corals’ thermal tolerance during ongoing climate change.

## Materials and Methods

### Sampling and thermal stress experiment

To study natural variation in thermal tolerance of corals, we measured responses to thermal stress in corals sampled from a natural population. For these experiments, we sampled 43 colonies of *Orbicella faveolata* from seven reef sites around the Bocas del Toro, Panama archipelago in 2015 (Figure 1a). Large intact colonies were extracted off the reef and tissue samples were collected and stored in RNAlater for genotyping (Scientific Permit No. SC/A-28-14). Each colony was cut into nine smaller uniform fragments (387 fragments total) with approximately 15-20 polyps per fragment. Fragments were maintained at ambient temperature in aquaria at the Smithsonian Tropical Research Institute (STRI) on Isla Colon, Bocas del Toro for one week prior to experimentation. Initial photographs of each individual fragment were taken before experiments began.

**Figure 1.**
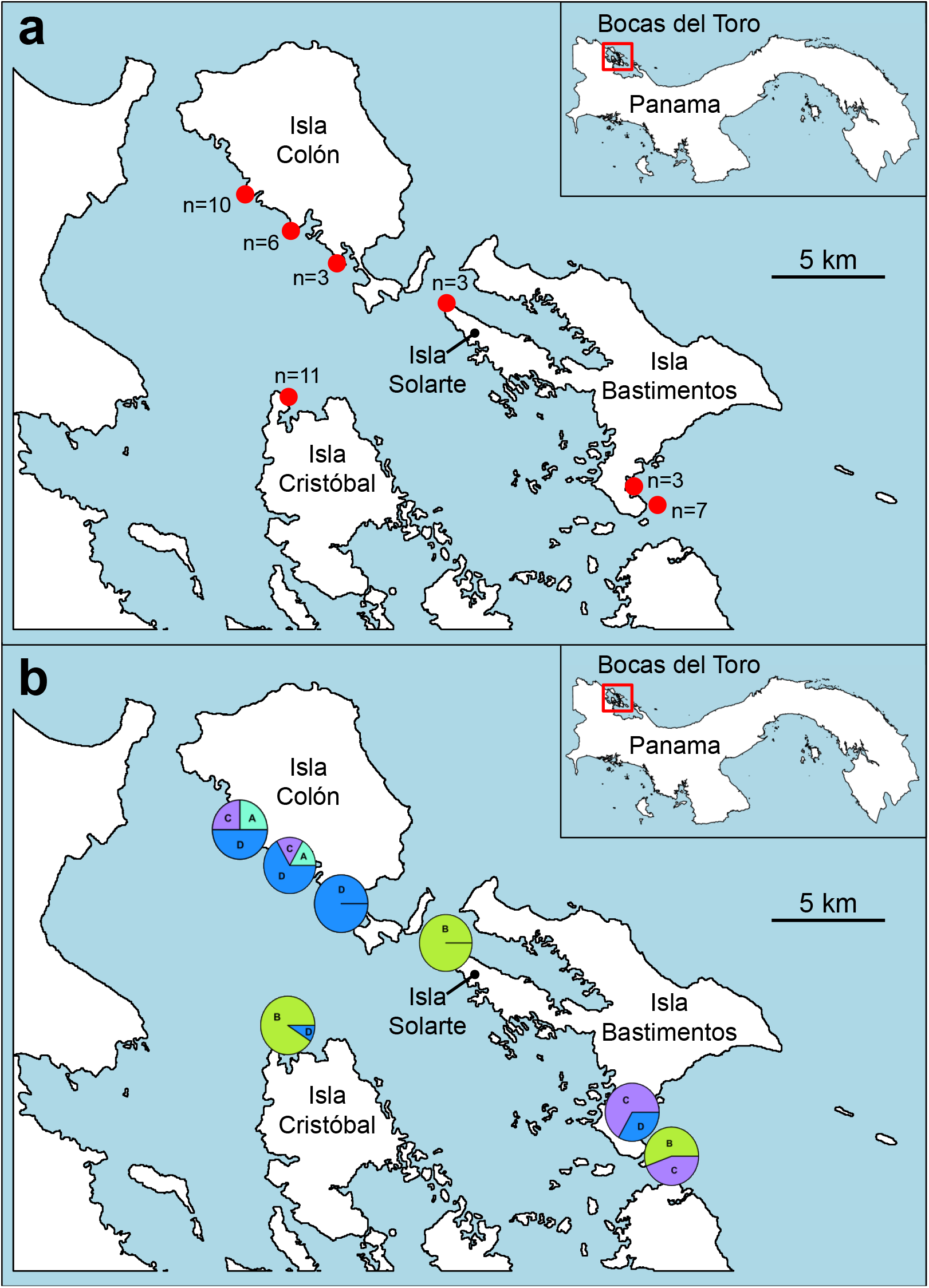
Map of Collection Sites around Bocas del Toro Archipelago, Panama. (a) Map of the seven locations where coral genotypes were collected around the archipelago. (b) Proportion of dominant symbiont types found at each site across colonies collected.

To estimate thermal tolerance, we exposed replicate fragments from each colony to a thermal stress treatment and measured their bleaching responses. Three randomly chosen fragments from each colony were maintained at control conditions (ambient seawater temperature of 29°C) while the remaining six fragments were ramped approximately 0.1°C every two hours to an elevated temperature treatment of 31°C for two weeks and 32°C for an additional two weeks. Corals were maintained for 4 weeks in normal and elevated temperatures, monitoring pH and salinity daily. Corals were monitored by daily visual inspection to evaluate bleaching response using the Coral Watch color scorecard, and the effects of temperature stress were scored as the number of degree heating weeks (DHW) required to induce bleaching. The experiment was terminated when approximately half of the fragments were bleached. Photographs were taken at the end of 4 weeks (approximately 5 DHW) and tissues were sampled and stored in RNAlater.

### Multilocus SNP genotyping of coral colonies

To estimate genetic relatedness and test for genetic associations with thermal tolerance, we conducted multilocus SNP genotyping on all coral colonies. To that end, we extracted genomic DNA from each colony using the Omega bio-tek E.Z.N.A. Tissue DNA Kit (Omega Bio-tek, Norcross, GA). We used the 2bRAD (Restriction Site-Associated DNA) protocol for SNP genotyping, a streamlined and cost-effective method for genome-wide SNP genotyping (Wang *et al.*, 2012). For these libraries we used the reduced tag representation method previously described (Wang *et al.*, 2012), using selective adaptor with overhangs ending in “NR” to target ¼ of the AlfI sites in the genome. This approach made it possible to analyze the number of samples included here on a limited budget, a tradeoff between marker number and sample numbers. We combined these libraries in equimolar amounts for sequencing in a single lane of 50 bp SE reads on Illumina HiSeq 3000 at OSU’s Center for Genome Research and Biocomputing (CGRB).

We analyzed the resulting data using a 2bRAD reference our research group has recently produced and used for a linkage map (Snelling *et al.*, 2017). Since the reference was produced from larval stages that naturally lack algal symbionts, no special filtering was required to eliminate algal reads in these samples from adult tissue. We conducted this analysis as previously described for de novo analysis of corals (Wang *et al.*, 2012; Howells *et al.*, 2016). Briefly, we filtered reads prior to analysis to exclude any low quality or uninformative reads (Joint Genome Institute, 1997), then aligned reads to the reference using SHRiMP (Rumble *et al.*, 2009) and called genotypes based on nucleotide frequencies at each position (calling loci homozygous if a second allele was present at less than 1%, heterozygous if present at > 25%, and leaving the genotype undetermined at intermediate frequencies where genotypes cannot be confidently determined from allele frequencies) (Wang *et al.*, 2012). Genotypes were called with a permissive threshold of ≥ 5x to call as many loci as possible for this genome wide survey of associations with bleaching responses. The scripts used for this analysis are available at (https://github.com/Eli-Meyer/2brad_utilities).

### Profiling algal symbionts with amplicon sequencing (ITS2)

To control for variation in the algal symbiont communities of each coral, which can contribute to variation in thermal tolerance of the holobiont (host plus associated algal and microbial symbionts) (Abrego *et al.*, 2008; Howells *et al.*, 2011), we sequenced the symbiont community in each colony using Sanger and Illumina amplicon sequencing. First, we amplified ITS2 using PCR primers previously described for studies of *Symbiodiniaceae* diversity (LaJeunesse, 2002), and sequenced the resulting amplicons using Sanger Sequencing. The resulting sequences were compared with multiple known ITS2 sequences from all formerly described *Symbiodinium* clades A-H (Hunter *et al.*, 2007; Cunning *et al.*, 2015). Using our symbiont sequences and these reference ITS2 sequences, we created an alignment in the program MEGA (Kumar *et al.*, 2017). A maximum likelihood phylogenetic tree was created with all known and unknown sequences to determine which clades our coral samples fell into. The dominant symbiont type was assigned for each sample by comparing the phylogenetic tree of unknown and known samples.

To confirm these results and evaluate whether our samples included mixed symbiont populations, we prepared additional ITS2 sequencing libraries for high-throughput sequencing on Illumina. We prepared these libraries using forward (5’-TACACGACGCTCTTCCGATCTGAATTGCAGAACTCCGTG-3’) and reverse (5’-ACGTGTGCTCTTCCGATCGGATCCATATGCTTAAGTTCAGCGGGT-3’) primers, and sequenced libraries using 300 bp PE read chemistry on Illumina MiSeq at OSU’s Center for Genome Research and Biocomputing (CGRB). We filtered reads to exclude any low quality reads (<20), removed reads lacking the expected amplicon primer sequence, and removed orphan reads. After filtering, paired reads from each sample were merged and were imported into dada2 (Callahan *et al.*, 2016). Using dada2, we further filtered samples for missing data and removed chimeric sequences. In this way, we identified valid amplicon sequence variants (ASVs) and described the abundance of each ASV in each sample. Finally, we created a BLAST database containing a diversity of annotated ITS2 sequences and the ITS2 sequence from the *Orbicella faveolata* host (Cunning *et al.*, 2015) and identified the clade of each ASV by comparison with this database.

We identified the dominant symbiont type in each colony based on the consensus of Sanger and Illumina sequencing results. While Sanger data lack resolution to describe mixtures of algal symbiont clade types, we interpret these sequences as the dominant symbiont types in each sample based on the presence of a single dominant haplotype in sequencing chromatograms. For Illumina, we quantified the proportion of each sequence variant in each sample and assigned a dominant clade if sequence variants were present >80% and a mix of symbionts if <80%. We included the dominant or mixed clade type(s) for each colony in quantitative models of bleaching responses to evaluate the contribution of variation in the dominant symbiont type to variation in thermal tolerance.

### Quantifying bleaching responses

To quantify bleaching in each fragment, we used qPCR to estimate the abundance of algal symbionts relative to host cells (Cunning *et al.*, 2015). We quantified collected samples after stress experiments in qPCR reactions. DNA from all fragments (control and heat-stressed from each colony) was extracted using an organic phase extraction. All qPCR reactions were run on an Eppendorf Realplex 4 machine using the SYBR and ROX filters. Each reaction consisted of 7.5 μL SensiFAST SYBR Hi-ROX master mix (Bioline, Taunton, MA), 4.3 μl NFW, 0.6 μl each of forward and reverse 10-μM primers, and 2 μl of genomic DNA (10ng total) in a final volume of 15 μl. The thermal profile for each reaction consisted of an initial denaturing step of 95°C for 2 min, followed by 40 cycles of: 95°C for 5 s, annealing temperature of 60°C for 30 s, and then 72°C for 30 sec. All control and heat-stressed samples were run using the same reaction parameters and were analyzed together. In addition, one sample was included on every plate as an inter-plate calibrator. We quantified host cells using host actin loci using the forward (5’-CGCTGACAGAATGCAGAAAGAA-3’) and reverse (5’-CACATCTGTTGGAAGGTGGACA-3’) primers, as previously described (Cunning *et al.*, 2015). To quantify *Symbiodiniaceae* in each sample we used a pair of universal primers developed based on multiple sequence alignments of the cp23S-rDNA locus from multiple *Symbiodiniaceae* species (https://www.auburn.edu/~santosr/sequencedatasets.htm). We identified regions that were sufficiently conserved to design primers suitable for qPCR (53-76 and 169-189 in that alignment). We conducted qPCR with primers (5’-CTACCTGCATGAAACATAGAACG-3’ and 5’-CCCTATAAAGCTTCATAGGG-3’) to determine the total amount of symbiont cells present after experimentation in control and experimental conditions. Host cell quantifications (C_T_ values) were subtracted from symbiont cell quantifications to calculate the dC_T_ value in each colony, a measure of the ratio of symbiont cells to host cells, for both control and experimental conditions. The dC_T_ stress value was subtracted from the dC_T_ control value to generate ddC_T_ values, representing the symbiont density. Then, we used these ddC_T_ values for each colony to calculate the fold change of symbiont abundance (2^−ddCt^), which were then log-transformed to compare across colonies. Additionally, we calculated the relative change within stress and control samples separately. The dC_T_ values from stress and control samples were calculated as described above, and these dC_T_ values were then compared to a reference control sample to generate ddC_T_ values, representing the symbiont density in all samples relative to a reference sample. We analyzed these qPCR data on relative symbiont density of each fragment to evaluate the effects of genotype, origin, and symbiont type on bleaching responses.

### Estimating heritability of variation in bleaching responses

Estimating the heritability of this variation in bleaching responses requires information on genetic relationships among subjects, which is initially unknown in samples collected from a natural population. For our study, we inferred genetic relatedness among samples based on multilocus SNP genotypes, and then used the genetic relatedness matrix derived from these SNPs to estimate genetic variance components. For this analysis we used the ‘related’ package in R and used the method described by Queller & Goodnight to calculate genetic distance between samples (Queller & Goodnight, 1989; Muir & Frasier, 2015; Tavalire *et al.*, 2018). After developing this matrix of genetic relatedness among samples, we analyzed variation in bleaching responses in the context of these relationships to estimate heritability. Using the R package ‘regress’, we created a linear mixed model with symbiont clade type and population source as fixed effects (site where samples were collected) as a fixed effect (Tavalire *et al.*, 2018). This analysis accounted for variation in thermal bleaching responses attributable to these specific factors. We estimated narrow-sense heritability and the associated standard error based on the phenotypic variation remaining after accounting for these known sources of variance, using the ‘h2G’ function in the R package ‘gap’ (Zhao, 2007).

### Testing for genetic associations with bleaching responses

To identify genetic markers associated with variation in bleaching responses, we tested for associations at each SNP locus using linear mixed models including SNP genotypes as a random effect and population source as a fixed effect. To account for errors arising from multiple tests, we converted controlled false discovery rate at 0.05 using the pFDR procedure (Storey, 2003). The multilocus SNP genotypes obtained from 2bRAD made it possible to test for associations between bleaching phenotypes and genotypes at each locus. Combining SNP data and the linkage map for this species (Snelling *et al.*, 2017), we searched for genomic regions underlying variation within more thermally tolerant phenotypes. We used the R package rrBLUP to test for associations between bleaching responses and genotypes at each locus, accounting for genetic structure in the population using an additive relationship matrix produced from SNP genotypes. We used the *A.mat* function to calculate the additive relationship matrix, considering all loci with no more than 5% missing data, then used the GWAS function in rrBLUP to conduct association tests, requiring allele frequencies > 0.08 (a second allele was detected at least 3 times), and included source population as a fixed effect. Once significant SNPs were found, we searched genomic scaffolds to examine neighboring genes. Based on an integrated genomic resource our group has recently developed by combining the linkage map with transcriptome and genome assemblies (Snelling *et al.*, 2017), we calculated linkage disequilibrium (LD) blocks in cM for each SNP based on <10% recombination frequency. We searched within each LD block to identify genes linked to each SNP.

### Profiling gene expression in heat-tolerant and susceptible colonies

To evaluate whether genomic regions associated with heat tolerance include genes differentially expressed between heat-tolerant and susceptible genotypes, we profiled transcriptional responses in a subset of corals demonstrating contrasting phenotypes (3 heat-tolerant, collected at Isla Bastimentos; 3 heat-susceptible collected at Isla Solarte) (Figure 2). RNA was extracted from replicate fragments from each colony using the Omega Bio-tek E.Z.N.A. Tissue RNA Kit (Omega Bio-tek, Norcross, GA). RNA was then used to prepare 3’ tag-based cDNA libraries for expression profiling (Meyer *et al.*, 2011). Samples were individually barcoded and combined in equal ratios for multiplex sequencing. We sequenced these libraries repeatedly on multiple runs because incompatibilities between the versions of the library preparation primers and the recently updated sequencing platforms resulted in very low sequencing yields. The first run was on the HiSeq 3000 platform at OSU’s CGRB, the second run on HiSeq 4000 at the University of Oregon’s Genomic and Cell Characterization Core Facility, and the third run on MiSeq at OSU’s CGRB. After sequencing, we processed the raw reads to remove non-template regions introduced during library preparation, and excluded reads with long homopolymer regions (>20bp) and low-quality reads with a Phred score of <30. All filtering steps were conducted using publicly available Perl scripts from https://github.com/Eli-Meyer/rnaseq_utilities. We mapped the high quality reads against the transcriptome for this species (Anderson *et al.*, 2016) using a short-read aligner software SHRiMP (Rumble *et al.*, 2009), and counted unique reads aligning to each gene to produce count data for statistical analysis of gene expression in each sample.

**Figure 2.**
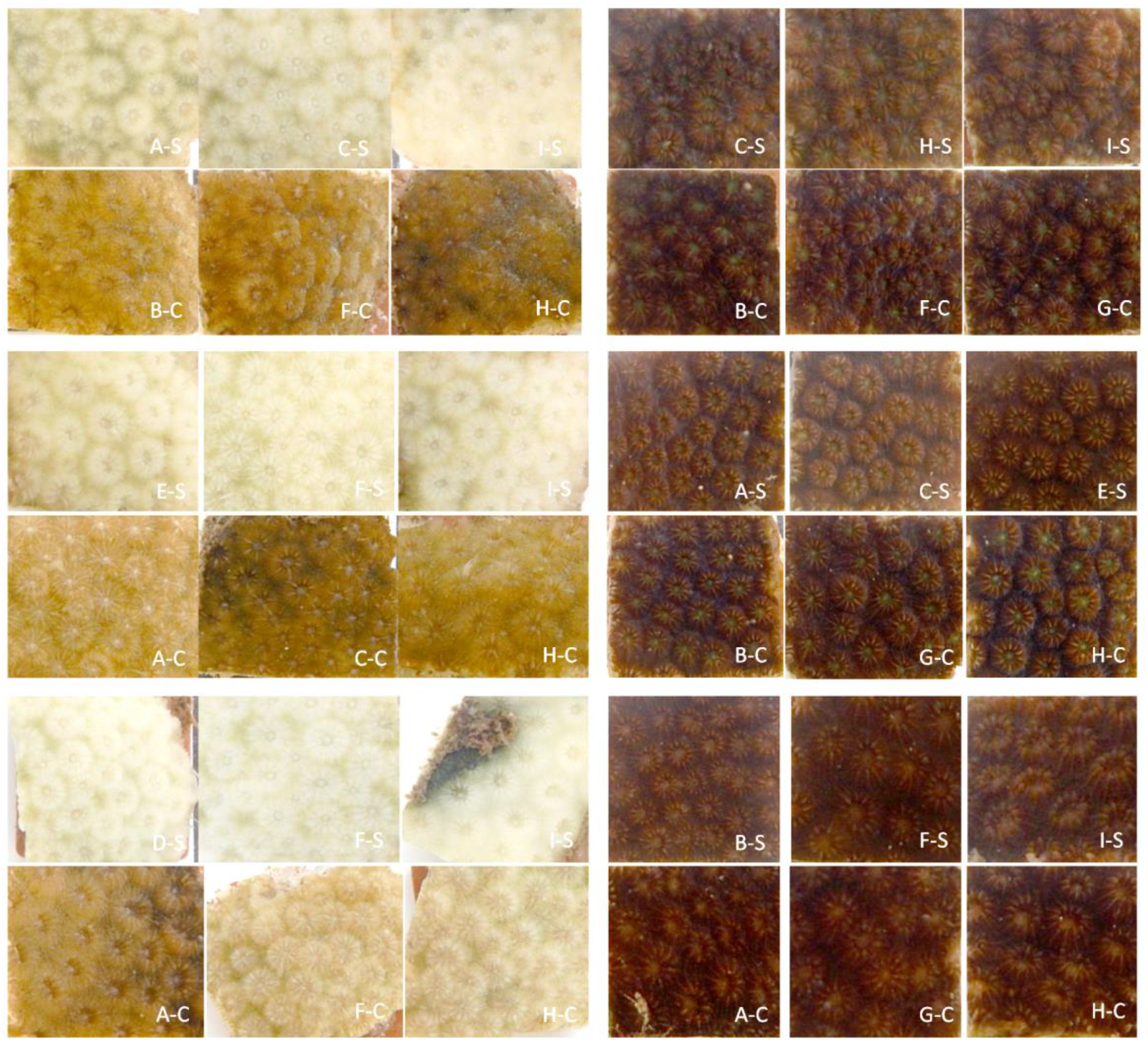
An example of the striking contrast between bleaching phenotypes of heat-susceptible and -tolerant corals sampled for this study. Each panel of six images represents fragments from a single colony, with control fragments indicated with “-C” (bottom of each panel) and heat-stressed fragments indicated with “-S” (top of each panel). Bleaching responses varied widely among colonies, but very little among fragments prepared from each colony.

We tested for differential gene expression using a negative binomial model in the R package DESeq2 (Love *et al.*, 2014). We tested for changes in gene expression by evaluating changes in stress-induced expression across samples in control and stress treatments. Our models tested for effects of treatment (control versus heat stress treatment) and bleaching response (susceptible versus tolerant) as main effects, and their interaction (treatment x response). We identified differentially expressed genes (DEGs) controlling false discovery rate at 0.05. To identify patterns of differential expression among the interaction effect DEGs, we conducted hierarchical clustering of expression patterns, subdividing the tree into clusters of correlated genes using the *cutree* function in R (Oksanen, 2010).

## Results

### Sequencing yield and SNP genotyping

To analyze genetic relationships among corals and associations with bleaching responses, we conducted multilocus SNP genotyping using a sequencing-based approach (2bRAD). Altogether, we sequenced 150 million high-quality reads, averaging 3.87 million reads per colony. We mapped these reads to a reference previously developed from aposymbiotic larvae, ensuring the loci being genotyped are derived from the coral host rather than the algal symbionts. We genotyped >700 kb at ≥ 5x coverage in each sample (Supplementary Table S1), identifying a large number of putative polymorphisms (35,067 loci). We further filtered genotypes to minimize missing data and genotyping errors, identifying a set of 5,468 high-quality SNPs that we used for all subsequent analyses.

### Symbiodiniaceae communities in host colonies and bleaching responses

To identify the dominant symbiont type or mixed symbiont communities in each coral colony, we sequenced ITS2 amplicons using Sanger and Illumina sequencing. In an effort to identify the dominant clade present in each colony, we classified the origin of each Sanger sequence by constructing a maximum likelihood tree including diverse representatives from *Symbiodiniaceae*, formerly described as *Symbiodinium* clades A-H (Hunter *et al.*, 2007; Cunning *et al.*, 2015). This analysis identified all sequences as members of clades A-D (Figure 1b; Supplementary Figure 1), and revealed differences in symbiont types across samples from different sites.

To confirm and expand on these results, we analyzed high-throughput ITS2 sequence data from the same samples. For this analysis we used a BLAST database with known ITS2 sequences (Cunning *et al.*, 2015) to classify the proportion of sequence variants in each sample originating from each symbiont species (or clade) (Figure 1b; Supplementary Figure 2). Most colonies contained a dominant sequence variant (>80%), while only two colonies showed a mixed community with two clade types. We considered both Sanger and Illumina data to assign the dominant or mixed symbiont community for each colony. Comparing these data revealed that the symbiont type identified from a single Sanger sequencing reaction in each sample corresponded to the dominant type identified in deep sequencing for nearly all samples in both datasets (26/29). For a small number of sample (3/29), the symbiont type identified from Sanger sequencing corresponded to a minor component of the community identified by Illumina sequencing rather than the dominant type. Illumina libraries for the remaining 14 samples were unsuccessful due to host contamination, so their identities were assigned based on Sanger sequencing. Overall, there was strong agreement between the assignments of symbiont type between Sanger and Illumina sequence data (Supplementary Figures 1, 2).

After 4 weeks in thermal stress at 31°C and 32°C, we saw considerable variation in bleaching among stressed fragments, while symbiont density changed very little across control samples (Figure 3). While there was variation between colonies, there was little to no variation in bleaching among fragments from the same colony (Figure 2). We quantified symbiont densities in each fragment using qPCR, and estimated the bleaching response of each colony as the log fold change between stressed and control fragments (Figure 3; Supplementary Figure 3). Colonies showed substantial variation in both their initial symbiont densities and their bleaching responses, based on both visual examination of the fragments and qPCR analysis of relative symbiont abundance (Figure 2 and 3). Most colonies bleached in response to thermal stress, but the extent of these bleaching responses varied considerably (Figure 3).

**Figure 3.**
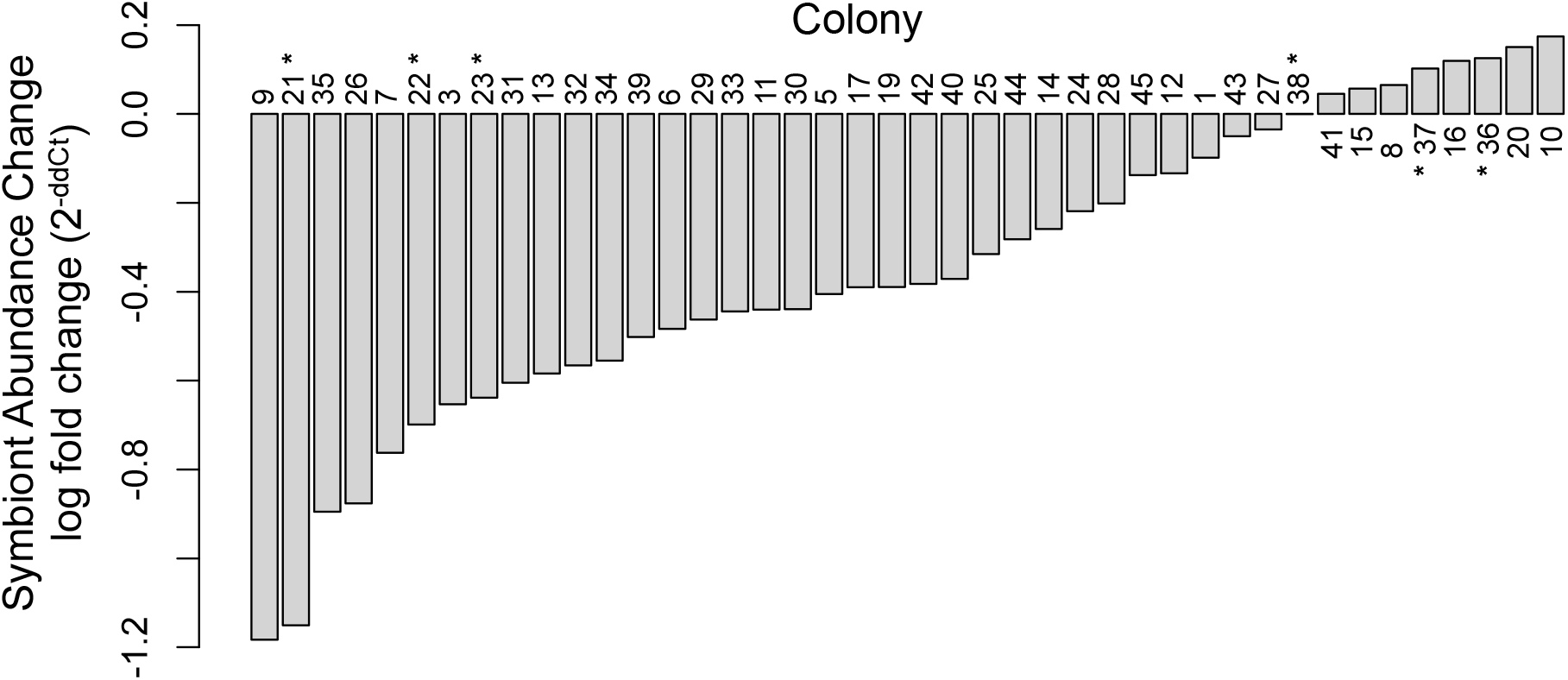
Quantification of algal symbiont densities using qPCR reveals variation in bleaching phenotypes. Bars represent the log fold change (2^−ddCt^) of symbiont abundance between control and stress samples across colonies after four weeks in control and experimental conditions. Starred colonies indicate those selected for RNASeq, showing contrasting abundances.

### Heritable variation in thermal tolerance in a natural population

To investigate heritable variation in thermal tolerance, we combined SNP data with bleaching responses measured by qPCR. We conducted a mixed model analysis to determine which factors to include in our heritability and association models. While population source had significant effects on thermal tolerance (*p*=0.0014), symbiont type had no effect (*p*=0.06). However, to be conservative, we included all factors in our REML mixed model to partition variation in thermal tolerance into genetic and non-genetic variance components. We estimated genetic relatedness among samples based on multilocus SNP genotypes, and then partitioned variance into genetic and non-genetic variance components in an ‘animal model’ (Wilson *et al.*, 2010) based on this genetic relatedness matrix. On average, pairwise genetic distances between colonies (calculated as the proportion of divergent alleles) between samples was 0.098 (range: 0.001 - 0.176). This analysis revealed that after accounting for effects of source and symbiont type, phenotypic variation in bleaching responses (log-fold change values) was highly heritable, with a narrow-sense heritability (*h^2^*) of 0.58 (SE=0.22). Taken alone, this estimate suggests substantial potential for adaptive responses to ocean warming in this population (but see Discussion for additional considerations).

### Genomic basis for variation in thermal tolerance

To understand the genomic basis for this variation in thermal tolerance, we used our SNP genotypes to test for associations between bleaching responses and genotypes. For this analysis, we conducted a series of linear mixed models testing for the effect of genotype at each locus while accounting for population structure. To visualize regions of the genome showing strong association with thermal tolerance, we mapped the results from statistical tests onto the integrated map, plotting −log_10_(p-value) for each marker by linkage group and position (Figure 4). After multiple test corrections, we found twelve markers significantly associated with bleaching; three markers when examining the fold change between control and stressed samples, three markers when examining the relative change in control samples, and six markers when examining the relative change in stressed samples (FDR ≤ 0.05). We emphasize that these three analyses of the symbiont densities are not intended to represent independent traits, but different aspects of biological variation relevant for thermal stress (initial symbiont density, bleaching response, and final symbiont density).

**Figure 4.**
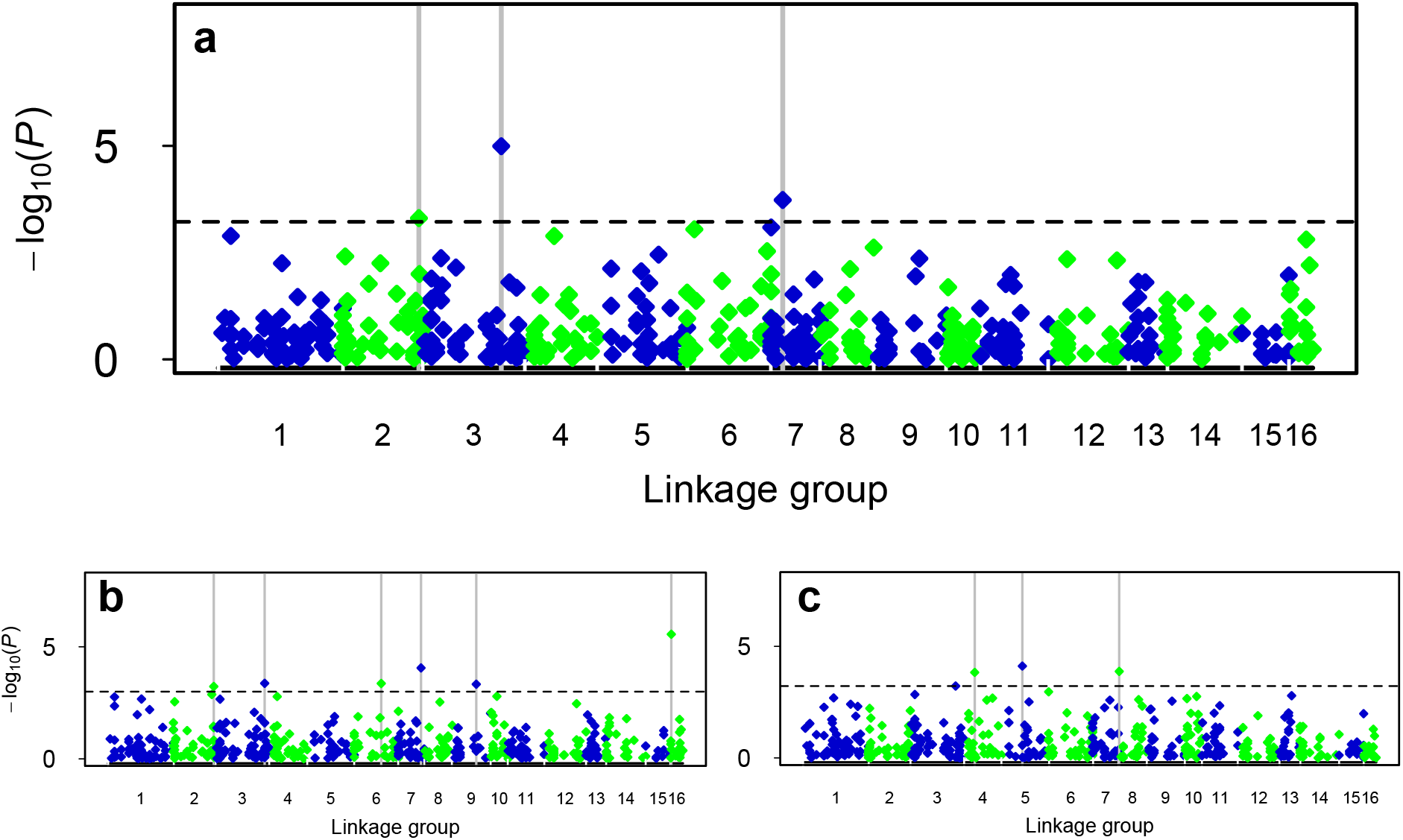
Mapping statistical associations between SNP genotypes and bleaching responses onto the linkage map identifies genomic regions associated with thermal tolerance in *O. faveolata* a) using the fold change between control and stress bleaching responses, b) relative change in stress samples, and c) relative change in control samples. Genetic markers are mapped against the linkage groups, indicated by alternating colors. Three markers on linkage groups 3, 5 and 7 were significantly associated (gray lines) with variation in bleaching responses during thermal stress, six markers on linkage groups 2, 3, 6, 7, 9 and 16 were significantly associated with variation in relative symbiont change in stress samples, and three markers on linkage groups 4, 5 and 8 were significantly associated with variation in relative symbiont change in control samples (FDR<0.05).

To identify the genomic positions of these SNPs were located and the genes linked to each marker, we used the integrated map (Snelling *et al.*, 2017) to search for genes closely linked (within an LD block) with each marker. Our SNPs fell onto linkage groups 2 (two SNPs), 3 (two SNPs), 4, 5, 6, 7 (two SNPs), 8, 9 and 16. Within the LD blocks around our SNPs, we used the integrated map to identify genes linked to each marker. All genes identified in this analysis are shown in Supplementary Table S2.

This analysis identified several groups of genes previously implicated in stress responses of corals or other Cnidarians, including genes with roles in oxidative stress responses, regulation of protein folding or degradation, and regulation of apoptosis. Genes linked to markers on LG2 included peroxiredoxin, a redox regulation protein for oxidative stress and genes associated with apoptosis (protein NLRC3). Genes linked to the markers on LG 3 were mucin proteins, ubiquitin protein ligases, caspase, a potassium voltage-gated channel protein, cellular tumor antigen p53, a gene associated with apoptosis; cytochrome 450, a protein involved in defense against chemical stressors; the chaperone DnaJ homolog involved in preventing inappropriate unfolding of proteins; heat shock protein 70, and glutathione s-transferase, a key enzyme in enhancing the oxidative stress response. Genes on LG 6 included ubiquitin protein ligases and potassium voltage-gated channel proteins. Catalase, apoptosis-inducing factor proteins, sodium-potassium transporting proteins involved in ion transport, tyrosine kinase receptor proteins involved in responding to oxidants and tumor necrosis receptor-associated proteins, important regulators of the apoptosis pathway were all linked to the markers on LG 7. Genes linked to the marker on LG 8 included mucin proteins and protein disulfide-isomerase, part of the unfolded protein response pathway. Genes linked to the markers on LG 9 and 16 included tyrosine kinase receptor proteins and tumor necrosis receptor-associated proteins, and ubiquitin protein ligases.

We also found several novel groups of genes that were not expected based on prior studies but were repeatedly observed across multiple markers and linkage groups in our study, suggesting a possible functional role for these genes in bleaching responses. These included 5-hydroxytryptamine (serotonin) receptors (5 genes altogether, linked to markers on LG 2, 3, 9, and 16). Similarly, we repeatedly found that galanin receptors were linked to bleaching associated markers (10 galanin receptor genes linked to bleaching-associated markers on LG 3, and 6). Galanins are neuropeptides classically associated with activities in the brain and peripheral nervous system, that have recently been shown to play diverse roles in innate immunity, inflammation, and energy metabolism (Lang *et al.*, 2014). We also found multiple collagen proteins (8 collagen genes linked to bleaching-associated markers on LG 2, 3, 6, 7, and 8). The possible functions of these genes in coral stress responses is not clear, but the repeated observation that these genes are linked to bleaching-associated markers on multiple scaffolds and linkage groups suggests that variation in these genes may contribute to variation in bleaching responses.

### Differences in transcriptional responses of tolerant and susceptible phenotypes

To further investigate the mechanisms of thermal tolerance, we profiled gene expression in contrasting phenotypes. For this dataset, we chose three heat-tolerant colonies and three susceptible colonies (Figure 2). The three heat-tolerant colonies were collected at Isla Bastimentos and contained clade C and D symbiont types, while the three heat-susceptible colonies were collected at Isla Solarte and all contained clade B symbionts (Figures 1, 2). These sites were approximately 15 km from one another and Isla Bastimentos exhibited more protection from wave action than Isla Solarte. Comparing bleaching responses, colonies from Isla Solarte had an average log-fold value of −0.8 (susceptible to bleaching) whereas colonies from Isla Bastimentos had an average value of 0.1 (tolerant to bleaching) (Figure 3).

Using a tag-based RNASeq approach (Meyer *et al.*, 2011), we prepared sequencing libraries for all 36 fragments (six colonies with six fragments, three control and three heat-stress fragments). We sequenced our libraries three times, once on Illumina HiSeq 3000, Illumina HiSeq 4000, and Illumina MiSeq, and all sequenced reads from all three runs were combined. In total, 63.9 million raw reads were produced, with approximately 1.73 million reads per sample. The majority of these passed quality and adaptor filtering (93%) leaving 59.4 million HQ reads for expression analysis (Supplementary Table S1).

Using a negative binomial model, we tested for changes in gene expression, evaluating differences in stress-induced expression. Our model tested for the effect of bleaching response, whether the colonies were bleached or unbleached, the effect of treatment, whether the fragments were in control or heat-stress, and the interaction effect between type and treatment. We found 737, 104, and 187 differentially expressed genes (DEGs) when testing for main effects of type and treatment, and their interaction, respectively. The interaction between type and treatment on gene expression can be visualized in a heatmap of expression for these DEGs (Figure 5), where heat tolerant colonies (red bars in figure 5) generally express these genes at higher levels than heat susceptible colonies (light blue bars in figure 5) regardless of treatment. Heatmaps for the effects of treatment and type are shown in Supplementary Figure 4. A complete list of differentially expressed genes in each category is provided in Supplementary Table S3.

**Figure 5.**
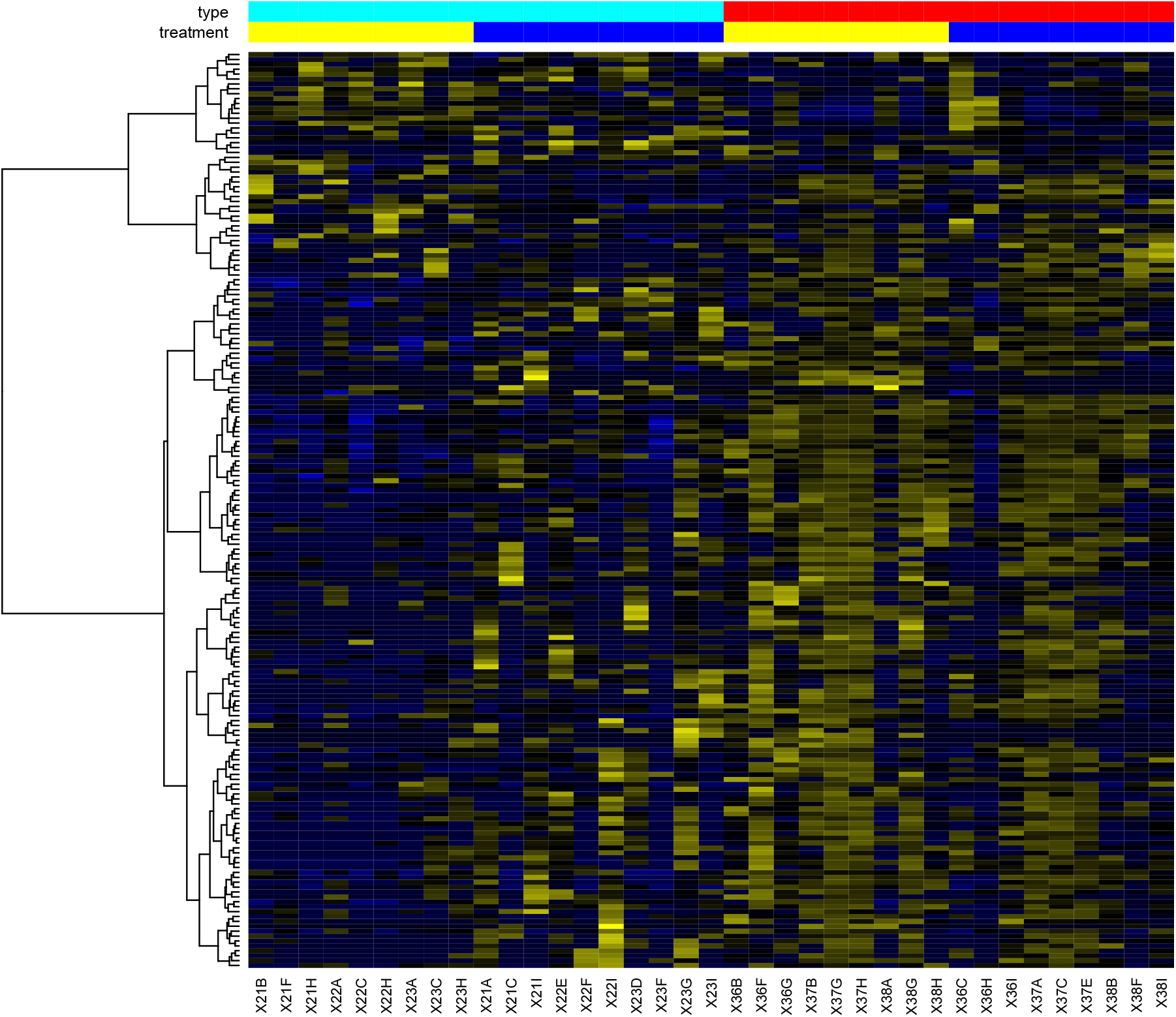
Heatmap showing relative expression of genes that were differentially expressed between heat-tolerant and heat-susceptible corals. The 187 DEG in this category are shown here, with samples and genes grouped with hierarchical clustering based on similarity in gene expression patterns. In the heatmap, blue indicates low expression, black moderate expression, and yellow indicates high expression. Colored bars indicate the type and treatment of each sample included in this analysis; red type refers to tolerant phenotypes, light blue refers to susceptible phenotypes, yellow bars are control samples, and dark blue bars are samples in heat stress treatment.

A substantial number of genes showed significant type × treatment effects, where the effects of treatment on expression differed between tolerant and susceptible corals. To characterize these interactions, we averaged expression for each gene in both susceptible and tolerant phenotypes for each treatment. Gene expression profiles were categorized into two dominant patterns. In the first patterns, genes were expressed at overall higher levels in heat-tolerant corals and were downregulated during thermal stress, and expressed at lower levels overall in heat-susceptible corals but upregulated during thermal stress. We found 159 genes in this category (Figure 6a). The second pattern was the opposite: genes that were expressed at higher levels overall and upregulated during thermal stress in heat-tolerant corals, and down-regulated in susceptible corals (Figure 6b). The remaining 33 genes formed a third cluster with similar patterns as 6b but with more variation across genes (not shown).

**Figure 6.**
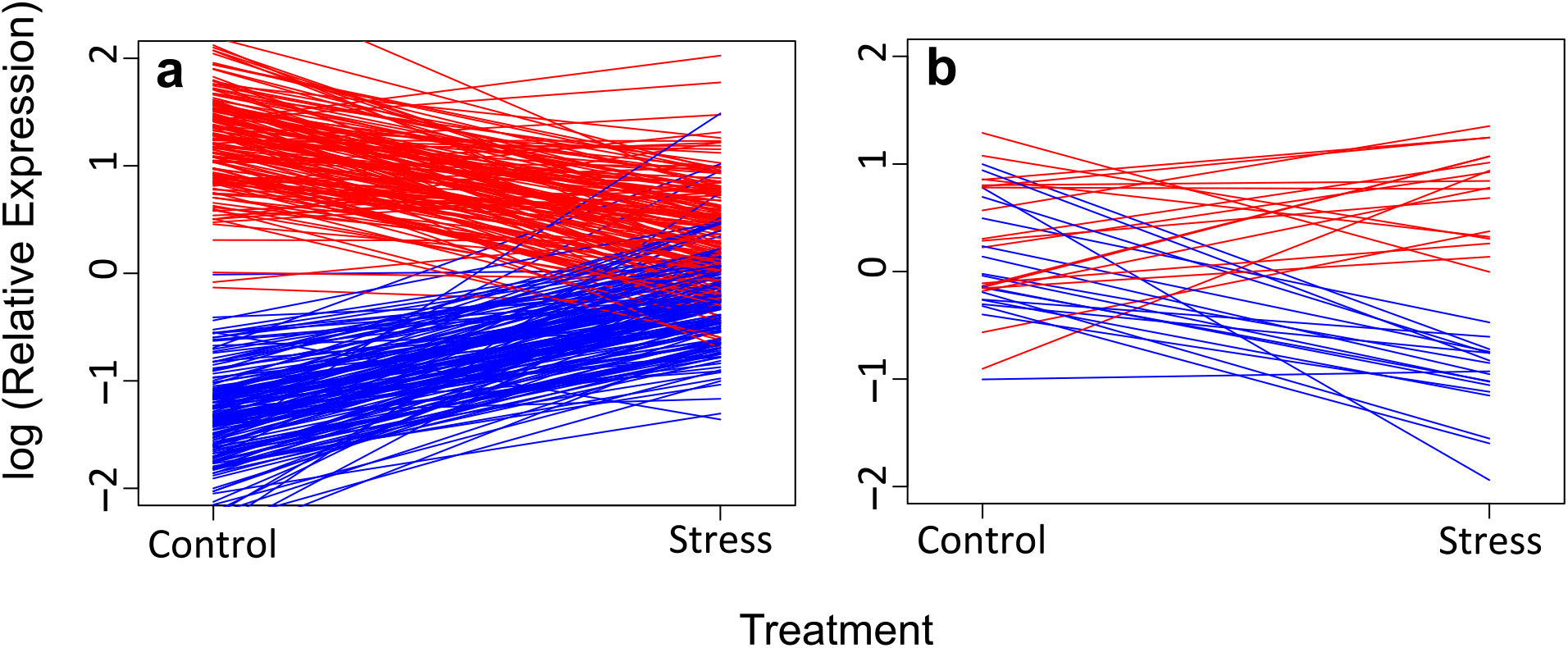
Type × treatment effects on gene expression fall into two general categories. a) 159 genes were downregulated in heat-tolerant corals (red) and upregulated in heat-susceptible corals (blue). b) 18 genes showed contrasting expression changes, but in the opposite directions: up-regulated in heat-tolerant corals and down-regulated in heat-susceptible corals. The remaining 33 genes showed similar patterns to (b) but were less consistent across genes, forming a third cluster (not shown).

Finally, we compared differentially expressed genes and those genes within the gene neighborhoods on our four linkage groups. Genes differentially expressed as a function of type, treatment or their interaction all contained ubiquitin protein ligases. In addition, when examining differentially expressed genes in the type effect, we found multiple collagen genes, mucins, as well as DnaJ proteins, glutathione peroxidase, and peptidyl-prolyl cis-trans isomerases, genes known to have a potential role in response to heat stress.

## Discussion

Our study provides some of the first quantitative estimates for heritability of variation in bleaching responses of corals. This builds upon larval studies (Meyer *et al.*, 2009a, 2011; Dixon *et al.*, 2015) that have demonstrated substantial heritability in responses to elevated temperatures, but left uncertainty in whether these findings extended to adult corals with intracellular algal symbionts and the energetic demands of calcification. Our findings confirm that some coral populations harbor similar genetic variation in thermal tolerance traits of adult coral colonies. These parameters have been studied in Indo-Pacific Acroporids, but to our knowledge no quantitative estimates for heritability of thermal tolerance were previously available for corals in the Robust clade (Fukami *et al.*, 2008; Meyer *et al.*, 2009a, 2011; Kitahara *et al.*, 2010; Baums *et al.*, 2013; Dixon *et al.*, 2015) or any other Caribbean corals. This is an important consideration because heritability of a trait is specific to the population and environment under study, suggesting caution in generalizing results from Indo-Pacific larval studies of Acroporids to evaluate potential adaptive responses in the deeply diverged groups of corals that dominate Caribbean reefs (Meyer *et al.*, 2009a, 2011; Baums *et al.*, 2013; Dixon *et al.*, 2015; Kenkel *et al.*, 2015; Lohr & Patterson, 2017).

To investigate the functional basis for this variation in bleaching responses, we conducted genomic and transcriptomic studies comparing allele frequencies and transcriptional stress responses in these corals. We found genetic markers significantly associated with thermal tolerance, and used the integrated genomic resource developed from a genetic linkage map and a genome sequence assembly to identify some of the genes linked to these markers. We found that transcriptional responses of heat-tolerant corals to thermal stress are markedly different from those of heat-susceptible colonies. We identified just under 200 genes differentially expressed as a function of type × treatment interactions, which were generally expressed at higher levels in tolerant corals and regulated in opposite directions by tolerant and susceptible corals in response to thermal stress.

This study builds on growing evidence that coral populations harbor genetic variation that may support adaptation to ocean warming. These questions are especially pressing for Caribbean corals, where reefs have declined severely over the last few decades (Hughes & Tanner, 2000). Since genetic variation supporting heritable variation in traits under selection is species- and population-specific, measuring these parameters in Caribbean populations is vital for understanding the future of these ecosystems. Our study documents considerable genetic variation in thermal tolerance for a population of the mountainous star coral, *Orbicella faveolata*, an important reef-builder throughout the Caribbean.

Our data suggests the genetic potential for substantial adaptive responses to selection for thermal tolerance in this population. Responses to selection can be modeled with the univariate breeder’s equation to estimate the expected rate of adaptation within a single generation (Falconer and Mackay, 1996). These predictions require empirical estimates for the narrow-sense heritability of the trait under selection, the proportion of phenotypic variation attributable to additive genetic variation (Falconer and Mackay, 1996). While it has been clear for some time that corals possess substantial variation in thermal tolerance, in part resulting from acclimatization or association with different algal symbionts (Fitt *et al.*, 2001; Howells *et al.*, 2011; Oliver & Palumbi, 2011; Silverstein *et al.*, 2012), the variation attributable to genetic factors in the coral host has remained understudied. This genetic variation will determine the adaptive responses of corals in the immediate future, since rapid adaptation relies on standing genetic variation in natural populations (Barrett & Schluter, 2008). Our study contributes novel information on this potential for adaptation to ocean warming, confirming that heritability of bleaching responses in adult corals can be comparable to the high heritability of thermal tolerance observed in some previous larval studies (Dixon *et al.*, 2015).

Importantly, these estimates of *h^2^* express genetic potential for adaptation, and other factors may constrain the adaptive responses that are actually realized in nature. The breeder’s equation expresses the rate of adaptive change within a single generation, requiring that we account for generation times to convert these estimates into units of adaptive change per decade or century. Massive corals like *Orbicella* are slow-growing and while direct estimates of generation time are unavailable for *O. faveolata* itself, comparisons with similar slow-growing massive corals suggests that these corals probably begin reproduction at ~ 5 years old and reach peak reproductive output around 10-15 years (Babcock, 1991). These life-history considerations impose inherent constraints on the rates of adaptation in this species, since even “rapid” adaptive changes occurring in a single generation would take 5-15 years to affect populations of adult corals. Additionally, correlations among traits can alter responses to selection relative to univariate predictions (Lande & Arnold, 1983; Houle, 1991; Falconer & Mackay, 1996; Lynch & Walsh, 1998). In these cases, selection for one trait affects the distribution of not only that trait, but also indirectly affects the distributions of correlated traits (Falconer & Mackay, 1996; Lynch & Walsh, 1998). Negative correlations among fitness related traits may constrain adaptive responses to selection (Etterson & Shaw, 2001), while positive correlations may facilitate adaptive responses (Agrawal & Stinchcombe, 2009). These correlations can change in different environments (Messina & Fry, 2003; Sgrò & Hoffmann, 2004), so describing these effects is also required for understanding responses to selection. Future studies should investigate the possibility that trait correlations may constrain adaptive responses in corals, preventing these populations from achieving the rapid adaptive responses that *h^2^* estimates suggest are possible.

The development of sequencing-based approaches for multilocus SNP genotyping has made genomewide association studies (GWAS) a widely used tool for identifying markers associated with traits of interest (Schlötterer *et al.*, 2015). These approaches map statistical associations between genetic markers and traits onto a genomic reference to identify regions of the genome underlying variation in the trait. Such an analysis obviously requires a genomic resource for mapping, and this requirement has limited the application of these approaches in many non-model systems. Despite limitations in the genomic resources available for *O. faveolata*, we used an integrated resource our group has recently established (Snelling *et al.*, 2017) to map statistical associations with bleaching responses onto the *O. faveolata* genome. It is important to acknowledge that our study was underpowered with only 43 genotypes (logistical constraints prevented us from further sampling, in this case). Despite the low power of our sampling design, we succeeded in identifying genetic markers associated with variation in bleaching responses. These likely represent the loci with the largest effects on thermal tolerance in these samples, with additional loci of smaller effects remaining undetected.

The markers we identified are linked to biologically interesting genes that could contribute to host thermal tolerance. For example, we found gene functions involved in oxidative stress, neural response, ubiquitination, protein folding regulation, and apoptosis. Glutathione s-transferase functions as an antioxidant enzyme in response to reactive oxygen species and has been shown to increase during thermal stress (Downs *et al.*, 2002; DeSalvo *et al.*, 2008; Polato *et al.*, 2010). Linkage with this gene could indicate an important role for oxidative stress response to reactive oxygen species (ROS) production during stress induced by increasing sea surface temperatures or pathogens (Downs *et al.*, 2002; DeSalvo *et al.*, 2008; Polato *et al.*, 2010). Voltage-gated proteins have also been characterized in *Nematostella vectensis* and have shown the importance of these proteins for maintaining cellular homeostasis, regulation of movement, and feeding. The process of ubiquitination labels proteins for degradation and expression of ubiquitin protein ligases may play an important role in increased tolerance to heat-stress (Finley *et al.*, 1987; Pickart, 2001; Welchman *et al.*, 2005; Shahsavarani *et al.*, 2012). For corals, these genes are highly correlated with increased thermal tolerance and are typically up-regulated in heat-stress corals with more damaged proteins (DeSalvo *et al.*, 2008; Barshis *et al.*, 2010; Lundgren *et al.*, 2013; Bay & Palumbi, 2015).

Some of the most interesting genes found are those involved in protein folding. The DnaJ chaperone plays an important role in the unfolded protein response (UPR) and is typically seen up regulated in response to elevated temperatures (≥ 32°C) in species such as *Acropora hyacinthus* (Ruiz-Jones & Palumbi, 2017) and *Stylophora pistillata, Porites sp.* and *Acropora eurystoma* (Maor-Landaw & Levy, 2016). This protein is also a co-chaperone of Hsp70, making it an important marker for thermal stress in corals (Cyr *et al.*, 1994; Walter & Ron, 2011). The close proximity (within a LD block of <10% recombination frequency) to our SNPs suggests a possible role for these genes in determining variation in thermal tolerance among colonies of *O. faveolata.*

We also found genes repeatedly observed across multiple markers and linkage groups, but their function and relationship to thermal tolerance in corals is unknown. These genes included galanin receptors, 5-hydroxytryptamine (serotonin) receptors, and collagen proteins. Galanin receptors are known to modulate neural responses and have been shown to play an important role in responses to stress, such as pain, emotional stimuli, and disease (Mitsukawa *et al.*, 2009; Lang *et al.*, 2014; Sciolino *et al.*, 2015). In cnidarians, 5-hydroxytryptamine (serotonin) receptors may serve as a neural signaling molecule and radiolabeling studies have localized the distribution of these proteins around nerve tissues within host cells (Hajj-Ali & Anctil, 1997; Dergham & Anctil, 1998; Westfall *et al.*, 2000). Collagen is a component of the extracellular matrix and may be related to wound healing and regeneration in cnidarians (Reitzel *et al.*, 2010; Stewart *et al.*, 2017). Despite these genes having an unknown role in thermal tolerance, their continued expression and linkage to significant SNPs indicate they may contribute to tolerance in the coral host.

In addition to genetic analysis, high-throughput sequencing has also enabled widespread application of RNA-Seq approaches to profile gene expression (Wang *et al.*, 2009). These methods have been widely adopted to study transcriptional responses to thermal stress in corals (DeSalvo *et al.*, 2008; Voolstra *et al.*, 2009; Leggat *et al.*, 2011; Meyer *et al.*, 2011; Oliver & Palumbi, 2011; Bellantuono *et al.*, 2012a, 2012b; Barshis *et al.*, 2013; Kenkel *et al.*, 2013; Palumbi *et al.*, 2014). One finding that has emerged consistently from these studies is the observation that corals vary widely in their transcriptional and phenotypic responses to thermal stress (Hunter, 1993; Ayre *et al.*, 1997; Marshall & Baird, 2000; Baums *et al.*, 2013). Many studies have demonstrated variation in gene expression among coral phenotypes, both in natural populations and in controlled studies (López-Maury *et al.*, 2008; DeSalvo *et al.*, 2010; Meyer *et al.*, 2011; Granados-Cifuentes *et al.*, 2013).

Here, we built upon these studies by quantifying variation in transcriptional responses to thermal stress in the context of known genetic relationships and thermal tolerance phenotypes. We found that heat-tolerant and -susceptible corals differed substantially in their responses to thermal stress. Focusing on the genes differentially expressed as a function of the type × treatment interaction, we identified a cluster of genes that were expressed at higher levels in heat-tolerant corals than their susceptible counterparts, and were down-regulated during thermal stress whereas susceptible corals up-regulated the same genes (Fig 6a). These included genes associated with protein metabolism (ribosomal protein genes, E3 ubiquitin protein ligase, and ubiquitin-conjugating enzyme E2), regulation of apoptosis (cathepsin-L and AP-1), and genes associated with calcium-binding (calcium-binding protein CML19, calretinin, neurocalcin, and a voltage-dependent L-type calcium channel).

Differentially expressed genes that overlapped with our association study included collagen genes, mucins, DnaJ proteins, and glutathione s-transferase. Inferring functional consequences from gene expression profiles is always uncertain, but these patterns suggest that thermal tolerance phenotypes in corals may be achieved in part by down-regulating energetically expensive processes such as protein synthesis, and in part by altering expression of the regulatory machinery controlling apoptosis.

We also identified a cluster of genes showing the opposite pattern (up-regulated by heat-tolerant corals during thermal stress), which included a fluorescent protein. These proteins are commonly reported in studies of Cnidarian stress responses (Smith-Keune & Dove, 2008; Rodriguez-Lanetty *et al.*, 2009; Roth & Deheyn, 2013), and our findings provide additional evidence these genes may play a role in variation among corals’ thermal tolerances.

Overall, our study provides a novel perspective on the potential for corals to adapt to ocean warming by estimating heritability of variation in thermal tolerance for a Caribbean reef-builder. We found that corals sampled from a natural population in Panama varied widely in their bleaching responses during an experimental thermal stress treatment. We used multilocus SNP genotyping to infer genetic relatedness among corals and estimate narrow-sense heritability (*h^2^*) for variation in bleaching responses, revealing that variation in this trait is primarily attributable to additive genetic variation. This suggests substantial genetic potential for adaptation to ocean warming in this population, although the complexities of multivariate selection suggest caution in predicting responses to selection from a single trait. We used the same SNP genotypes to test for associations between bleaching responses and genotypes at each marker, identifying genetic markers for bleaching responses that can be directly applied in restoration and conservation efforts to identify heat-tolerant corals. We used expression profiling to demonstrate that heat-tolerant corals respond to thermal stress differently than susceptible corals, and functional analysis of the differentially expressed genes suggests differential regulation of protein metabolism and apoptosis in heat-tolerant corals. Our findings provide crucial data for models aiming to predict corals’ adaptation to ocean warming, and identify genetic markers for thermal tolerance that may be useful for restoration efforts as conservation biologists work to reverse the global degradation of coral populations resulting from changing ocean conditions.

## Acknowledgements

The authors are grateful to the Smithsonian Tropical Research Institute and the Institute for Tropical Ecology and Conservation for use of their facilities and for assistance with sample collection and fieldwork logistics. We thank Rachel Vasta and Chelsea Eckhart for assistance with sample processing. We thank Oregon State University’s College of Science for travel assistance through a travel grant to KD.

## Supplementary Information

Supplementary Table S1. Excel spreadsheet. Summary of sequencing yields, processing, and mapping efficiencies for 2bRAD and RNASeq sequencing libraries.

Supplementary Table S2. Excel spreadsheet. Complete list of genes linked to SNP markers significantly associated with variation in bleaching responses (within 10 cM of the markers).

Supplementary Table S3. Excel spreadsheet. Complete list of genes differentially expressed as a function of temperature, thermal tolerance type, or their interactions.

Supplementary Figure 1. Sanger sequence data results for all samples (colonies).

Supplementary Figure 2. Illumina MiSeq sequence data results for all samples (colonies).

Supplementary Figure 3. Quantification of algal symbiont densities using qPCR reveals variation in bleaching phenotypes as a function of symbiont types and site.

Supplementary Figure 4. Heatmaps showing relative expression of genes that were differentially expressed between heat-tolerant and heat-susceptible corals and between heat-stressed and control samples.

## Data Accessibility

Reference numbers for data in public repositories: sequence data archived at NCBI’s Sequence.

Read Archive under accession SUB3108676.

Scripts used for analysis can be found at https://github.com/Eli-Meyer.

## Authors’ Contributions

KD and EM conceived the study and conducted analyses. KD collected samples, conducted thermal stress experiments, prepared sequencing libraries, and wrote the manuscript. HE contributed to sample collection and fieldwork assistance. HT developed the heritability analysis pipeline. All authors have read and approved the manuscript.

