## Supplementary Figures for "Heritable variation in bleaching responses and its functional genomic basis in reef-building corals (*Orbicella faveolata*)"

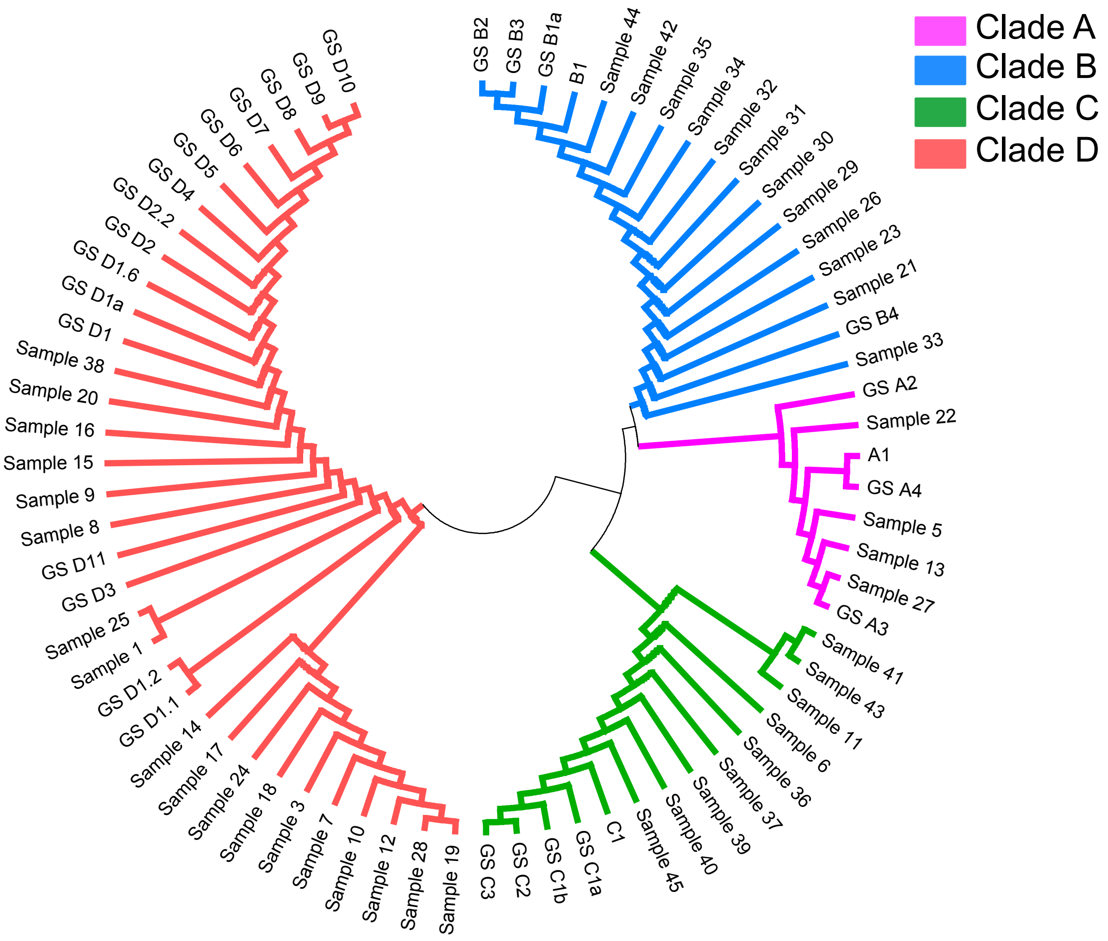


**Supplementary Figure 1.** Sanger sequence data results for all samples (colonies). Sanger sequence data resolved a dominant clade, A-D for all samples.

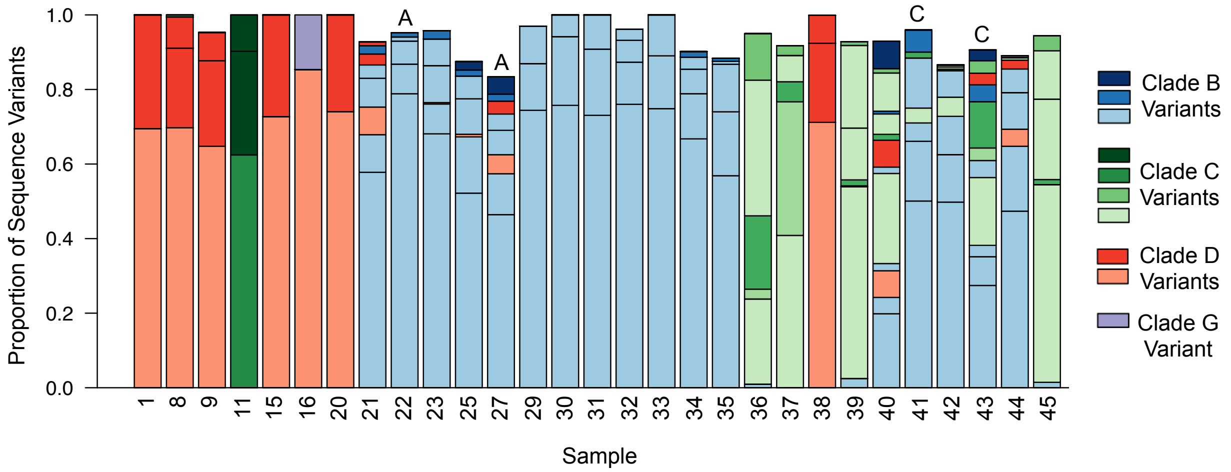


**Supplementary Figure 2.** Illumina MiSeq sequence data results for all samples (colonies). Symbiont variants (Clades B, C, D and G) are shown as a proportion within each sample. Letter above a column indicates samples that demonstrate disagreement between Sanger result and ITS amplicon result when assigning dominant symbiont clade.

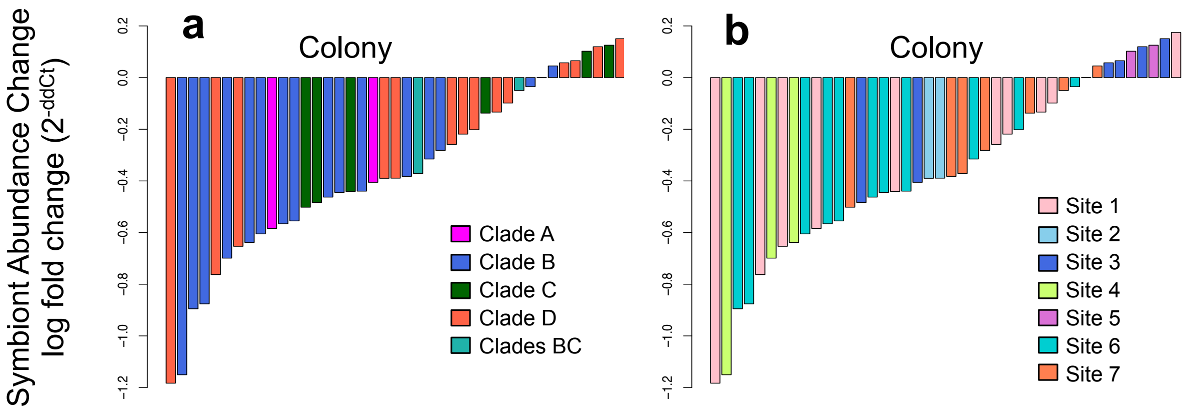


**Supplementary Figure 3.** Quantification of algal symbiont densities using qPCR reveals variation in bleaching phenotypes as a function of a) symbiont types and b) site. Bars represent the log fold change ($2^{-ddCt}$) of symbiont abundance between control and stress samples across colonies after four weeks in control and experimental conditions.

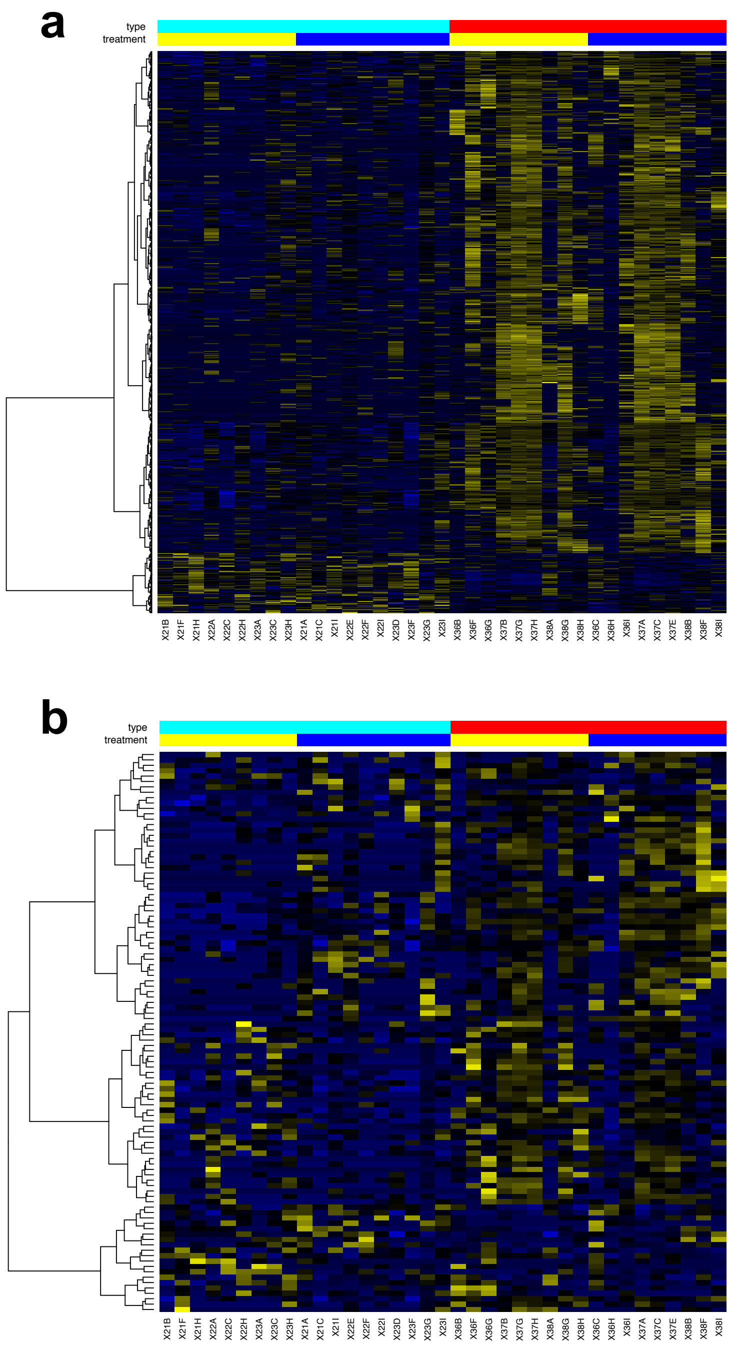


**Supplementary Figure 4.** Heatmap showing relative expression of genes that were differentially expressed **a)** between heat-tolerant and heat-susceptible corals (type effect; 737 DEGs) and **b)** between heat stressed and control samples (treatment effect; 104 DEGs). In both heatmaps, blue indicates low expression, black moderate expression, and yellow indicates high expression. Colored bars indicate the type and treatment of each sample included in this analysis; red type refers to tolerant phenotypes, light blue refers to susceptible phenotypes, yellow bars are control samples, and dark blue bars are samples in heat stress treatment.
